# Genetic Diversity and Phylogenetic Relationships of Annual and Perennial *Glycine* Species

**DOI:** 10.1101/557439

**Authors:** Eun-Young Hwang, He Wei, Steven G. Schroeder, Edward W. Fickus, Charles V. Quigley, Patrick Elia, Larissa Costa, Susan Araya, Marcio Elias Ferreira, Perry B. Cregan, Qijian Song

## Abstract

We estimated average genetic diversity of two *Glycine* annual and six perennial species based upon 76 orthologous gene sets and performed phylogenetic analysis, divergence analysis and tests for departure from neutrality of the eight species using 52 orthologous gene sets. In addition, 367 orthologous gene sets were used to estimate the relationships of 11 *G. canescens* accessions. Among the perennials, *G. canescens* showed the highest nucleotide diversity and the other perennials except *G. tomentella* had higher nucleotide diversity than the two annuals. Phylogenetic analysis of the *Glycine* showed a similar genome grouping with the previous report except *G. cyrtoloba* and *G. stenophita* formed a sister clade in the study. Divergence analysis supported the phylogenetic relationships, *G. falcata* was the most divergent from *G. max*, followed by *G. cyrtoloba*, *G. syndetika*, *G. tomentella* D3, *G. stenophita* and *G. canescens*. Neutrality selection tests within species showed that most genes were subjected to a recent directional selection due to a selective sweep or rapid population expansion. Although most gene sequence had negative and significant Tajima’s D, the sequences were homogeneous in the levels of polymorphism and divergence between *G. max* and other *Glycine* species based on the HKA test, thus, *Glycine* perennials may have experienced very similar evolutionary selection as inferred by trans-specific mutation analysis. The greater genetic diversity of most perennial *Glycine* species and their origins from the warmer and drier climates of Australia suggested the perennials as potential sources of heat and drought resistance that will be of value in the face of climate change.

## INTRODUCTION

The genus *Glycine* Willd. includes two subgenera, *Glycine* and *Soja* (Moench) F.J. Herm. Subgenus *Glycine* contains 25 named and at least six unnamed perennial taxa (Gonzalez-Orozco et al. 2012; Sherman-Broyles et al. 2014) and subgenus *Soja* contains two annual species, *Glycine max* (L.) Merr., the cultivated soybean, and *Glycine soja* Sieb. & Zucc., the wild soybean. The two subgenera shared a common ancestor approximately 5 million years ago (Innes et al. 2008; Egan and Doyle 2010), however, the biogeography of *Glycine* is unusual in that the annuals are native to northeastern Asia whereas the diploid (2n=38, 40) perennials are almost exclusively Australian (Doyle et al. 2004).

The perennial *Glycine* species have been demonstrated to be potential sources of economically important traits, especially disease resistance genes, for use in cultivated soybean. Approximately 41% of 294 accessions of 12 perennial *Glycine* species examined had moderate or high resistance to soybean rust (Hartman et al. 1992) and *G. canescens* had at least four soybean rust resistance loci (Burdon 1988). In addition, some accessions of most perennial *Glycine* species showed resistance to one of the most destructive soybean pests, the soybean cyst nematode (SCN) (Bauer et al. 2007; Riggs et al. 1998). These authors also determined that progenies derived from *G. max* (SCN susceptible) x *G. tomentella* (SCN resistant) showed resistance to SCN. These studies verified the perennial *Glycine* as a source of useful genes to improve cultivated soybean although currently hybridization between annual and perennial *Glycine* species is limited (Ratnaparkhe et al. 2011). The other important aspect of the perennial *Glycine* is that some of these species have been adapted to harsh environments, especially very dry and hot areas.

Early studies were focused on the collection of germplasm and the classification of the collection based upon crossability testing, cytogenetic analysis, as well as molecular marker analysis such as isozyme grouping and nucleotide variation in nuclear or chloroplast DNA (Singh and Hymowitz 1985a; Brown et al. 1990; Singh et al. 1992; Kollipara et al. 1994, 1997; Singh et al. 1998; Doyle et al. 1999a; Doyle et al. 2003b; Grant et al. 1984; Doyle et al. 1999b; Doyle et al. 2003a; Ratnaparkhe et al. 2011; Sherman-Broyles et al. 2014). These efforts established the phylogenetic relationships among the *Glycine* species indicating that the two annuals belong to the G genome group and the 25 perennials (Doyle et al. 2004; Sherman-Broyles et al. 2014) to the nine genome groups from A to I (Hymowitz et al. 1998). However, because of the lack of DNA sequence information for the perennials, relatively little is known about the genetic diversity of the perennial *Glycine* species and the relationships among the perennial species as well as their relationship with cultivated soybean. As the relationship could be an indicator of the successful rate of the crossing and the difference between the perennial species and cultivated soybean, the information could help us to choose perennial species for the improvement of cultivated soybean.

The development of high throughput DNA sequence analysis makes it possible to identify many orthologous gene sets in the *Glycine* species with a high efficiency. In this study, we estimated the average genetic diversity of six perennial *Glycine* species chosen to represent the phylogenetic diversity of subgenus *Glycine* (Ratnaparkhe et al. 2011), *G. canescens*, *G. cyrtoloba*, *G. falcata*, *G. stenophita*, *G. syndetika*, and *G. tomentella* D3, and the two annual *Glycine* species, *G. soja* and *G. max*, based upon the DNA sequence analysis of 76 orthologous gene sets. The phylogenetic analysis, evolutionary divergence analysis among the species and test for departure from neutrality were performed using common sequences of 52 orthologous gene sets with 87.6 Kbp in length to infer the relationships among the 77 accessions of the *Glycine* species. In addition, approximately 462 Kbp of sequences of 367 orthologous gene sets were used to estimate the relationships of 11 *G. canescens* accessions.

## MATERIALS AND METHODS

### Plant materials

Both annual *Glycine* species, *G. soja* (G genome) and *G. max* (G genome), were used for this study along with six perennial *Glycine* species, *G. canescens* (A genome), *G. cyrtoloba* (C genome), *G. falcata* (F genome), *G. stenophita* (B’ genome), *G. syndetika* (A genome; previously named *G. tomentella* D4), and *G. tomentella* D3 (D genome), chosen to represent the major clades of subgenus *Glycine* (Sherman-Broyles et al. 2014), which are well-correlated with the “genome groups” classified by Singh and Hymowitz (1985a) and Hymowitz *et al*. (1998).

There were two sets of accessions of each species used for the current study. The data set used for the estimation of genetic diversity was based on 100 accessions and the data set used for a phylogenetic analysis, evolutionary divergence analysis among the species and test for departure from neutrality included 77 accessions of the eight *Glycine* species. For the additional phylogenetic analysis of *G. canescens* 11 accessions were used that include 9 accessions used for the phylogenetic analysis of the *Glycine* species and two additional accessions. The accessions used for each study are listed in Supplementary File 1. The perennial *Glycine* species are basically inbred, they have similar low out-crossing rates as cultivated soybean. In the greenhouse, the pods from most species were formed even though the flowers were not blossom. In addition, very few heterozygotic single nucleotide polymorphic sites were observed from our sequences of the gene sets within each species.

### Sequence diversity analysis

#### Genomic DNA extraction

Genomic DNA of 10 to 15 accessions of each species (Supplementary File 1) was isolated from young leaves of greenhouse-grown plants using the CTAB method (Keim et al. 1988). All accessions were selected based on geographical locations that covered the entire range over which each species had been collected in an effort to sample the breadth of diversity within each species (Figure 1).

**Figure 1.**
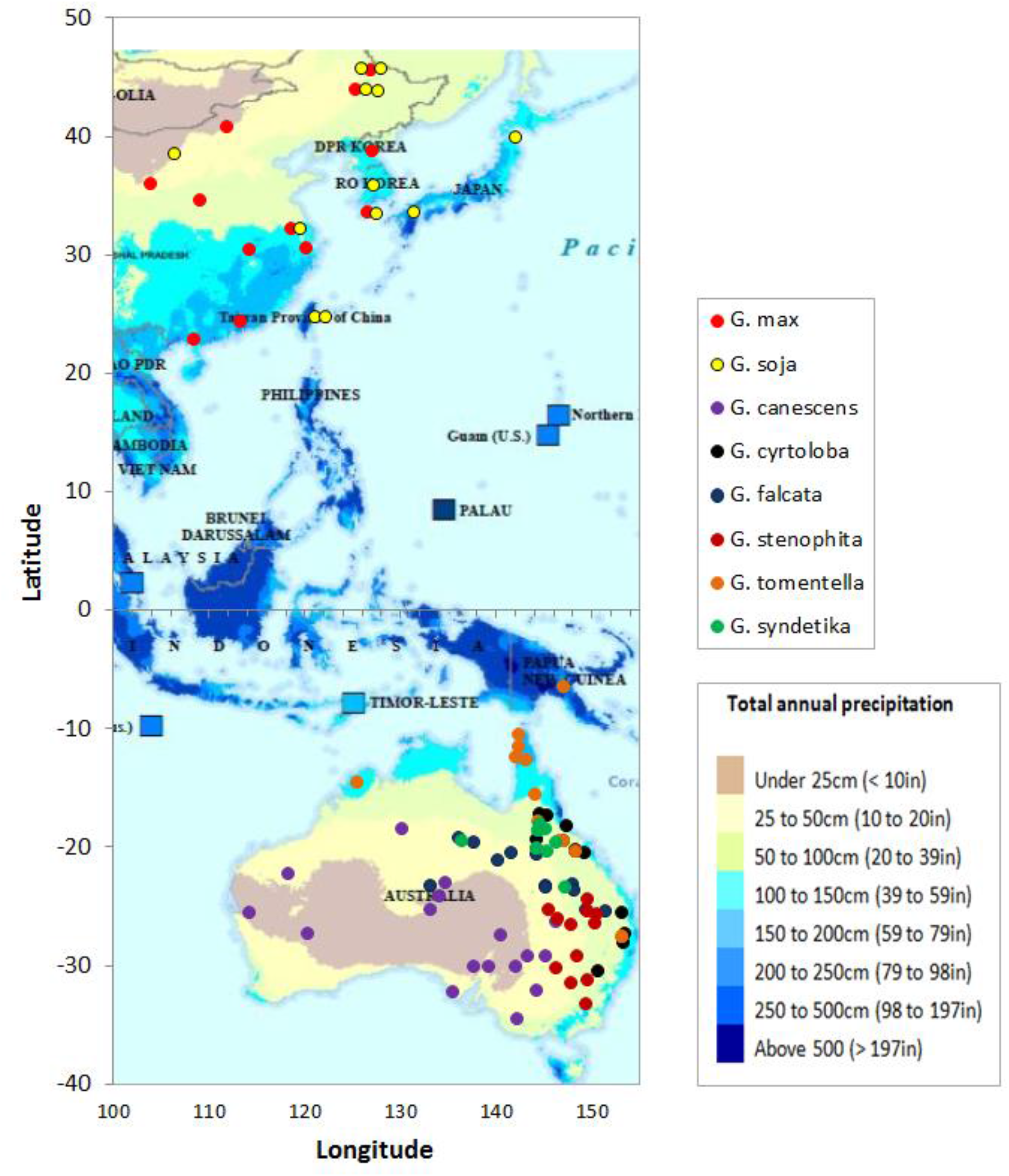
Geographical origin of the accessions. The origin of 98 accessions of the two annual and the six perennial *Glycine* species and the average annual precipitation in the corresponding regions.

#### RNA extraction and cDNA preparation

A seed of one accession from each of six perennial species was scarified and planted in a 1:1 soil and sand mix and grown in the greenhouse. Leaves, pods, and flowers were harvested from the single plant and roots were harvested from 5 to 10 seedlings grown on a wet paper towel in a petri-dish for 3 to 7 days. Total RNA was extracted from leaf, pod, flower and root tissue of each of the six perennial species using the Qiagen mini-RNA prep (Qiagen, Valencia, CA). Poly-A+ RNA was isolated from the total RNA using a Dynabead mRNA purification kit (Invitrogen, Carlsbad, CA) and cDNA libraries were constructed following the protocol “Preparing samples for sequencing of mRNA” (Illumina Inc., San Diego, CA).

#### Sample preparation for sequence analysis

The 6 cDNA libraries constructed from different tissues of the six perennial species were fragmentased for 20 min using NEB Next dsDNA fragmentase (NEB, Beverly, MA), end repaired and an ‘A’ overhang was added to the ends of the fragments. The end repaired cDNA libraries were ligated with the Illumina paired-end sequencing multiplex adapters and run on a 2% agarose gel for size selection. Fragments ranging from 100 to 150 bp and from 450 to 550 bp were isolated and normalized to reduce the relative number of highly expressed genes (Zhulidov et al. 2004) using double strand specific nuclease (Evrogen, Moscow, Russia). The normalized cDNA was PCR amplified for 17 or 18 cycles. The concentration of the amplified cDNA was verified using the Agilent 2100 Bioanalyzer (Agilent Technologies, Palo Alto, CA). The cDNAs from the different tissues of the same accession were combined to obtain equimolar concentrations of each species and run on the Illumina Genome Analyzer IIx to obtain 105 bp paired-end sequence reads.

#### Sequence analysis and selection of orthologs

The Illumina Off-Line Basecaller V1.8 software was used for base calling and demultiplexing. The reads from each species were assembled using Velvet software (v.1.013) (Zerbino and Birney 2008). The number of assembled contigs of one accession from each of the six species with sequence length greater than 400 bp and 1,000 bp ranged from 18,971 to 27,057 and 1,589 to 3,550, respectively. The resulting scaffolds from each species were blasted to each of the other perennial species using reciprocal BLAST at an E-value threshold of e-20 and a total of 295 cDNA sequence scaffolds (putatively orthologous gene sets) were identified that were present in each of the six species. These scaffolds were subsequently aligned to the Williams 82 soybean whole genome sequence (http://phytozome.jgi.doe.gov/pz/portal.html#!info?alias=Org_Gmax) to identify the corresponding orthologs in *G. max* and *G. soja*.

#### Species-specific primer design, PCR amplification, and sequence analysis

A total of 295 sets of species-specific PCR primer pairs were designed with an average predicted amplicon length of 1 Kbp using Primer3 (Rozen and Skaletsky 2000). In the case of the annual species, *G. soja* and *G. max*, only one set of 295 primer pairs was selected based upon the Williams 82 soybean genome sequence (Schmutz et al. 2010). The primers were used to amplify genomic DNA of one or two genotypes from each species. The resulting PCR products were run on a 2% agarose gel to determine if a single amplicon was produced. In those cases when a single amplicon appeared to be produced, the resulting PCR product was sequenced on an ABI3130xL (Applied Biosystems, Foster City, CA). To verify that a single amplicon was present, the resulting sequence was analyzed using the DNA analysis software Phred (Ewing et al. 1998) and Phrap (http://www.phrap.org/) and the resulting alignments and trace data were visually inspected in the Consed viewer (Gordon *et al*. 1998) to distinguish, as suggested by Choi *et al*. (2007), those amplicons that were locus specific and those that apparently resulted from amplification of two or more paralogous loci. Finally, of the 295 primer sets designed to putatively orthologous genes, 76 sets that produced a single amplicon in at least six of the eight species were selected (Supplementary File 2) and used for the amplification of genomic DNA of the 10 to 15 accessions from each of the eight species. The resulting amplicons were run on agarose gels and the concentrations were visually compared. Based on these relative concentrations, approximately equimolar amounts of the amplicons from the same accession were combined and the concentration of the combined DNA was determined using the Agilent 2100 Bioanalyzer. The DNA pool of the amplicons from each accession was prepared for sequence analysis on the Illumina HiSeq 2000 to obtain 150 bp single-end reads following the protocol described above, except, 96 sets of indexing adaptors were used for multiplex sequencing which were selected from the adaptors reported by Craig *et al*. (2008) and Hamady *et al*. (2008) and modified to allow paired-end sequence analysis using the Illumina HiSeq 2000. The resulting paired-end reads were de-multiplexed using CASAVA (v. 1.82) and assembled using Velvet software.

#### SNP and indel identification and the estimation of nucleotide diversity

The scaffolds of each accession were clustered using the 76 gene DNA sequences of their respective species obtained by Sanger sequencing on an ABI3130xL using USEARCH (Edgar 2010). The clustered sequences were aligned using MUSCLE with the gap open penalty of −400, gap extension penalty of 0, center parameter of 0, number of iterations of 16 and distance measure for initial iterations of Kmer4-6 (Edgar 2004). If any portion of an amplicon was missing from an accession, PCR was done using the species-specific primers and the PCR product was sequenced on an ABI3130xL. The sequence was analyzed using Phred and Phrap, after which the sequences were combined with the previously clustered scaffold sequences for SNP and indel detection. Nucleotide diversity, *θ* (Watterson 1975) and *π* (Tajima 1983), of each species was calculated to estimate the genetic diversity of the eight species.

### Phylogenetic analysis

#### RNA extraction, cDNA preparation, and sample preparation for sequence analysis

A seed of 10 to 12 accessions of the eight *Glycine* species was scarified and germinated on wet paper in a petri-dish for 3 to 7 days. The germinated seedling was transferred on top of a paper wick in a plastic pouch filled with nutrient solution (Barker et al. 2006) and grown for 3 to 14 days. Total RNA was extracted from a seedling of 10 to 12 accessions of each of the eight *Glycine* species from which poly-A+ RNA was isolated. cDNA libraries were constructed following the protocol “Preparing samples for sequencing of mRNA” from Illumina Inc. and prepared using the 96 sets of indexing adaptors following the protocol described above, except, DNA fragments ranging from 250 to 450 bp were isolated. The prepared cDNA libraries from the whole seedlings of the accessions of the same species were normalized, combined and run in a single lane on the Illumina HiSeq 2000 to obtain 150 bp paired-end sequence reads.

#### Sequence analysis and the selection of orthologs

The number of assembled contigs of 10 to 12 accessions of each of the eight species with a length greater than 400 bp ranged from 18,070 to 47,579. A total of 153,106 *Phaseolus vulgaris* L. and 189,593 *Vigna unguiculata* (L.) Walp ESTs were downloaded from NCBI and of these, 90,364 of the *P. vulgaris* and 125,808 of the *V. unguiculata* ESTs with sequence length greater than 400 bp were used for the selection of orthologs to serve as out-groups in the phylogenetic analysis of the *Glycine* species. To identify orthologs among the eight *Glycine* species and the two out-groups, cDNA sequences from the 95 *Glycine* accessions and ESTs of *P. vulgaris* and *V. unguiculata* were analyzed at an e-value threshold of e-10 using Proteinortho v 5.11 (Lechner et al. 2011). Proteinortho identifies a single ortholog as well as co-orthologs in those cases where there was more than one gene with very similar sequences (paralogs) found in an accession. If there were co-orthologs present in any accession of the eight species, the entire orthologous set was eliminated. In addition, it was difficult to obtain orthologous genes present in all 95 accessions so if the ortholog was present in greater than three accessions in each species, the gene was selected for the analysis. The resulting number of selected orthologous gene sets was 52 which were present in 77 of the 95 accessions of the eight *Glycine* species. To verify that the same homeologous gene sets were being compared for the phylogenetic analysis, evolutionary divergence analysis among the species and test for departure from neutrality, the selected 52 gene sets from each of the perennial species were aligned against the whole transcriptome sequences of the corresponding species using BLAST at an e-value threshold of e-20. The results showed that the second best alignment score of the 52 gene sets across the six species averaged only 19.8% of the best alignment score and the highest of the secondary alignment scores in any of the six perennial species was 82.5%. In addition, in the case of *G. max* and *G. soja*, we examined the potential homeologs of the 52 genes in *G. max* based on alignment to the whole genome sequence of *G. max* in an effort to further identify paralogous sequences. The mean alignment score of the 2nd best alignment averaged only 26.9% of the best alignment score. The best of the 52 secondary alignment scores was only 76% of the corresponding first alignment score. This was further indication that the 52 genes very likely have only one copy in the genome. Therefore, the 52 orthologous gene sets were used for the phylogenetic analysis, the evolutionary divergence analysis among the species and the test for departure from neutrality.

For a separate phylogenetic analysis of *G. canescens*, the assembled contigs and scaffolds of 11 accessions including the nine accessions used for the phylogenetic analysis of the eight *Glycine* species and two additional accessions whose transcriptome sequences were missing a number of the 52 genes used in the analysis of the eight species, and ESTs of the two out-groups, *P. vulgaris* and *V. unguiculata*, with a length greater than 400 bp were selected. Proteinortho was used to identify orthologs among the 11 *G. canescens* accessions. A total of 367 gene sets that did not include any of the 52 genes used in the analysis of the eight *Glycine* species were present in at least 7 of the 11 accessions and were used to estimate the relationships among the *G. canescens* accessions.

#### Phylogenetic analysis procedures

The 52 orthologous gene sets identified in the 77 accessions of the eight *Glycine* species and the 367 gene sets of the 11 *G. canescens* accessions were analyzed using MUSCLE (Edgar 2004) and concatenated using linux commands. The concatenated sequences were analyzed with 1,624 models using jModelTest 2.1.3 (Darriba et al. 2012) to select the best nucleotide substitution model. The 1,624 models included 203 different partitions of the Generalised time-reversible (GTR) rate matrix combined with rate variation (+I, +G, +I+G) and equal/unequal base frequencies. The likeliscore for the number of models for the best-fit models within the full set of 1,624 models was calculated and optimized at most 288 models while maintaining model selection accuracy. The best model selected for the orthologs of the *Glycine* species and the 11 G*. canescens* accessions was the same, TPM1uf+I+G. The best models selected by jModelTest 2.1.3. were used in Phyml v3.0 (Guindon et al. 2010) with 100 bootstrap replicates. The phylogenetic relationships of the *G. canescens* accessions were used to produce a phylogeography of the accessions using GenGIS (Parks et al. 2013). Digital map data were provided by the ORNL DAAC spatial data access tool (http://webmap.ornl.gov/).

### Analysis of evolutionary divergence of genic sequences and identification of trans-specific polymorphisms among *Glycine* species

Evolutionary divergence among *Glycine* species based on the aligned sequence of 52 genes was estimated using the Tajima-Nei distance method assuming equal substitution rates among sites and between transitional and transversional substitutions (Tajima and Nei 1984).

The distance matrix was calculated using software Mega 7 (Kumar et al. 2016). Trans-specific polymorphisms among species were also determined based on the aligned sequence of 52 genes using software Mega 7.

### Test of gene for departure from neutrality

Tajima’s D test was used to determine the selection neutrality of 52 genic sequences within species. Under the neutrality selection model, the scaled difference of the mean number of pairwise differences and the number of segregating sites of the sequences is expected to be the same (Tajima 1989). Because the theoretical distribution of Tajima’s D for 95% confidence interval is between −2 and +2 (Carlson et al. 2005), the D value greater than 2 or less than −2 was used as thresholds to determine significance of departure from neutrality selection. Calculation of Tajima’s D statistics was performed using software Mega 7.0 based on the aligned sequence for each gene, we choose the site sequence coverage above 70% as a threshold to eliminate the sites with missing bases. In order to test the correlation between the levels of within- and between-population DNA variation as predicted by the neutrality, equilibrium and independence and to test heterogeneity in levels of polymorphism relative to divergence (Haddrill et al. 2005), HKA tests (Hudson et al. 1987) of the genes between *G. max* and each of other *Glycine* species were performed using the DNASP Beta Version: 6.12.01 downloaded at http://www.ub.edu/dnasp/.

## RESULTS

### Genetic diversity of the *Glycine* species

A total of 76 gene sets were selected for the estimation of genetic diversity in the *Glycine* species. The number of fragments amplified in 10 to 15 accessions of each species that produced a single amplicon varied from 63 to 74 and the total length of sequence analyzed ranged from 37.9 to 45.0 Kbp in the eight species (Table 1). The nucleotide diversity measured as *θ* of the two annual species was 0.0011 and 0.0022 for *G. max* and *G. soja*, respectively, and that of the perennial species had up to four times higher genetic diversity than *G. max* with the highest nucleotide diversity in *G. canescens* (*θ*=0.0043bp) followed by *G. stenophita*, *G. cyrtoloba*, *G. syndetika*, *G. falcata* and *G. tomentella* D3 (Table 1). An average nucleotide diversity estimated as *π* also showed a similar trend with the diversity measured as *θ* where *G. canescens* had the highest diversity (*π*=0.0031bp). However, *G. stenophita* and *G. falcata* had relatively low *π* values compared with their *θ* values. The orthologous gene sets used for this study were distributed on 19 of the 20 soybean chromosomes (Figure 2) and included more than 37 Kbp of genic sequence in each of the eight species.

### Phylogenetic analysis of the *Glycine* species and *G. canescens*

**Figure 2.**
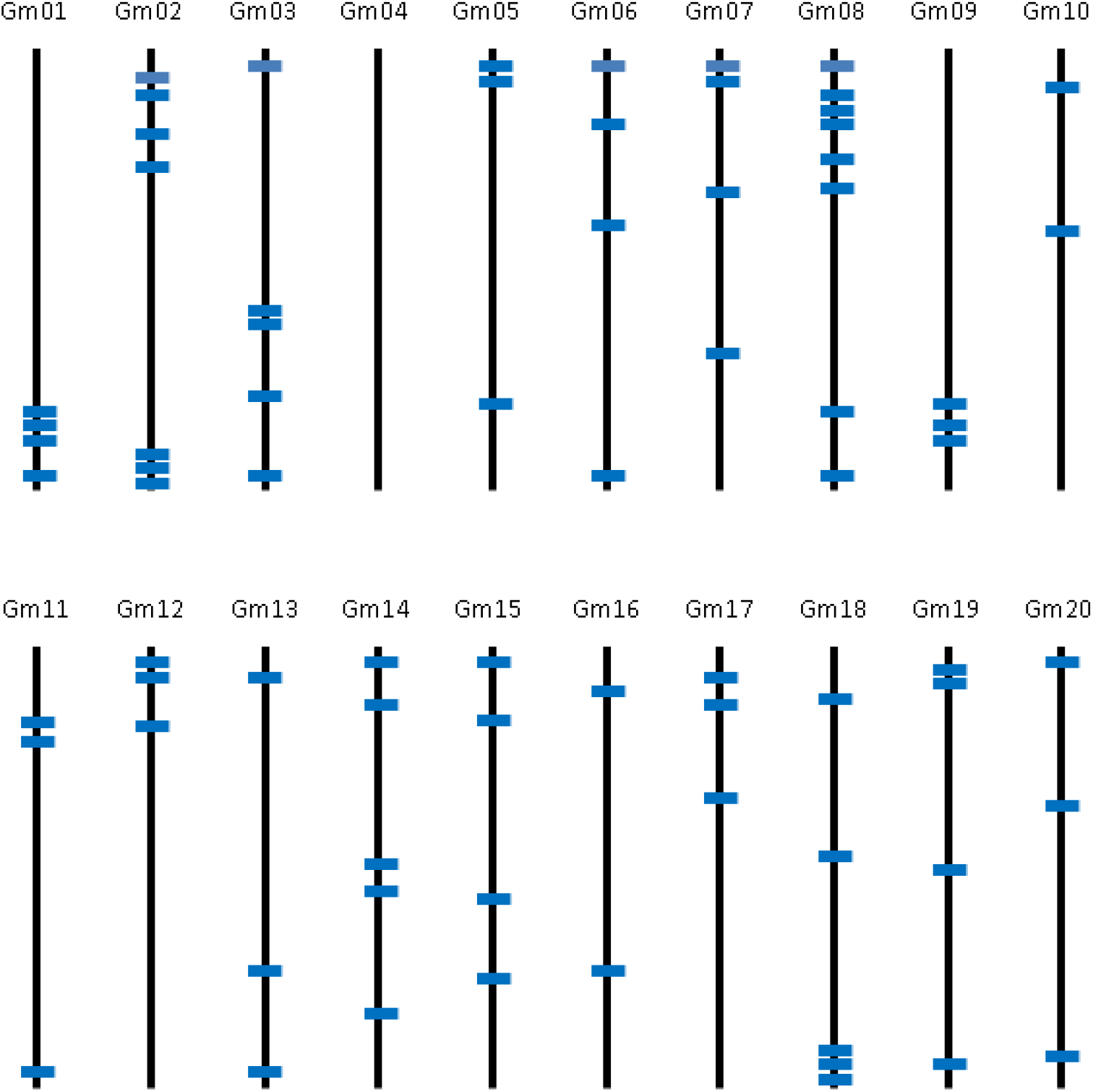
The locations of 76 orthologous genes on the 20 *G. max* chromosomes (Gm1-Gm20). The bars indicate the positions of the 76 genes used for the genetic diversity analysis.

**Table 1.** Nucleotide diversity (ϴ and π) of the Glycine species based upon the DNA sequence analysis of orthologous genes identified from 76 orthologous gene sets that produced a single amplicon in at least six of the eight Glycine species analyzed using species specific PCR primers.

| Species | Number of individuals | Number of genes | Total length (bp) | Number of SNPs | Nucleotide diversity ($\theta$) | Nucleotide diversity ($\pi$) |
| --- | --- | --- | --- | --- | --- | --- |
| *G. max* | 15 | 70 | 41,179 | 148 | 0.0011 | 0.0009 |
| *G. soja* | 12 | 70 | 45,044 | 295 | 0.0022 | 0.0016 |
| *G. canescens* | 15 | 74 | 44,560 | 626 | 0.0043 | 0.0031 |
| *G. cyrtoloba* | 12 | 63 | 37,931 | 366 | 0.0032 | 0.0029 |
| *G. falcata* | 12 | 65 | 37,904 | 307 | 0.0027 | 0.0016 |
| *G. stenophita* | 12 | 71 | 41,276 | 406 | 0.0033 | 0.0019 |
| *G. syndetika* | 10 | 69 | 42,120 | 365 | 0.0031 | 0.0025 |
| *G. tomentella* D3 | 12 | 67 | 41,684 | 202 | 0.0016 | 0.0010 |

A total of 52 orthologous gene sets present in 77 accessions of the eight *Glycine* species and two out-groups, *Phaseolus vulgaris* L. and *Vigna unguiculata* (L.) Walp, produced a concatenated data set approximately 87.6 Kbp in length. The phylogenetic analysis produced a phylogenetic tree (Figure 3) whose topology was quite similar to the genome grouping based upon Histone H3-D (Doyle et al. 1996; Sherman-Broyles et al. 2014), except *G. cyrtoloba* and *G. stenophita* formed a sister clade in this study. The annuals, *G. soja* and *G. max* of the G genome group, were strongly supported as a monophyletic group sister to the perennials. Within the perennial species, *G. falcata* was a sister of all the other perennial species with a bootstrap value of 67. As expected, the two A group species, *G. canescens* and *G. syndetika*, formed a sister clade which was a sister of the D group species, *G. tomentella* with a bootstrap value of 100. *G. cyrtoloba* of the C group and *G. stenophita* of the B’ group formed a clade which was a sister of the clade including the two A group species, *G. canescens* and *G. syndetika*, and the D group species, *G. tomentella*. The relationships of the clades were supported with moderate and high bootstrap values.

**Figure 3.**
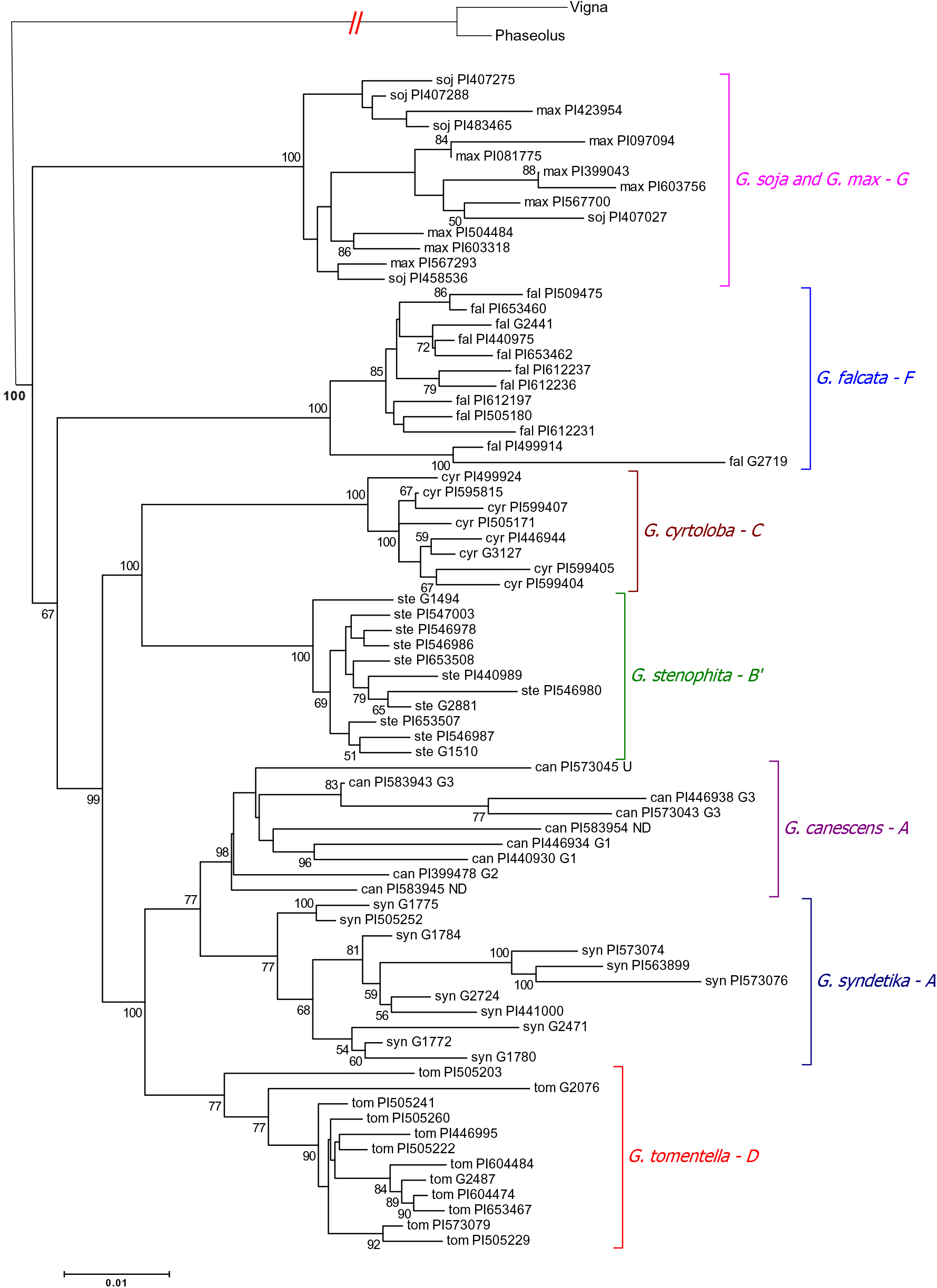
Phylogenetic trees of the *Glycine* species. The phylogenic tree of the 77 accessions of the eight *Glycine* species is based upon 52 orthologous gene sets. The letter next to each species indicates its nuclear genome designation as defined by Singh and Hymowitz (1985), Hymowitz (1998), and Doyle (2004b). The designation following the PI numbers of the *G. canescens* accessions indicates the subgroup designation (G1 through G3 and U = unassigned to a subgroup) assigned by Brown *et al*. (1990). *G. canescens* accessions followed by “ND” were not included in the Brown *et al*. (1990) study and thus, their subgroup was not determined. Nodes are based on 100 bootstrap replicates. The bootstrap values lower than 50 were eliminated. *Phaseolus vulgaris* L. and *Vigna unguiculata* (L.) Walp are used as out-groups.

Of the perennial *Glycine* species, *G. canescens* was the only species whose genetic structure was estimated based on isozyme variation by Brown *et al*. (1990). Thus, the PI number of each *G. canescens* accession is followed by the group (G1, G2, G3, and U: the heterogeneous, ungrouped accession) as assigned by Brown *et al*. (1990) (Figure 3 and Figure 4). In Figure 3, the relationships among the nine *G. canescens* accessions showed that there were two clades, one with the group G3 accessions and the other with the group G1 and one of ND accessions, which were sisters of the group G2, U and one of ND accessions. To verify the relationships among the *G. canescens* accessions in Figure 3 we performed a separate phylogenetic analysis using 367 orthologous gene sets identified in the 11 *G. canescens* accessions and the two out-groups which produced concatenated sequences with a length of 462 Kbp. The results showed that there were two major clades, one included accessions from the genome group G3 and the other included the remainder of the accessions. The latter clade included two clades, the first with the U and ND accessions and the second from the group G1 accessions, as well as the single G2 accession (Figure 4A). In addition, the phylogeography of the *G. canescens* accessions showed that the distance between the geographic origins of the accessions increased as their phylogenic relationships were more distant (Figure 4B).

**Figure 4.**
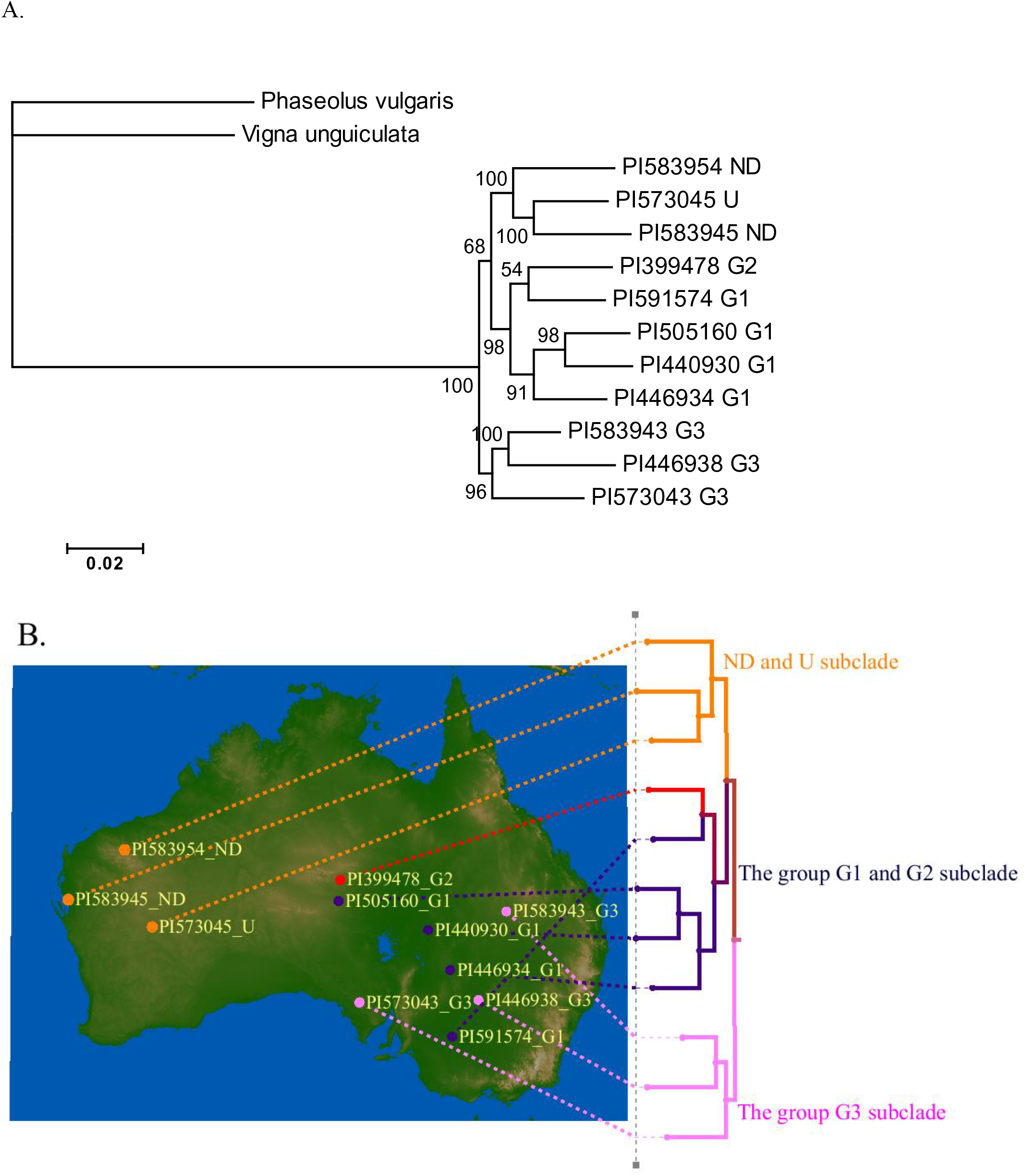
Phylogenetic tree and phylogeography of *G. canescens*. A) The phylogenetic tree is based upon 367 gene sets in the 11 accessions. The letter next to the PI numbers indicates the subgroup designation assigned by Brown *et al*. (1990). *G. canescens* accessions followed by “ND” were not included in the Brown *et al*. (1990) study and thus, their subgroup was not determined. Nodes are based on 100 bootstrap replicates. *Phaseolus vulgaris* L. and *Vignaunguiculata* (L.) Walp are used as out-groups. B) The geographic origin of each accession of the phylogeny is indicated as a dot on the map and the accessions defined as the same genome group by Brown *et al*. (1990) are assigned the same color. The PI number and the genome group of each accession is indicated next to the dot and the three subclades are indicated.

### Divergence among species

The Tajima-Nei distance analysis based on the 52 gene sequences (Supplementary File 3) showed that the divergence between *G. max* and *G. soja* is the smallest (0.009), perennial species *G. falcata* was the most divergent from *G. max*, followed by *G. cyrtoloba*, *G. syndetika*, *G. tomentella* D3, *G. stenophita* and *G. canescens*. Among the perennials, the largest differentiation was observed between *G. falcata* and *G. cyrtoloba*, and between *G. falcata* and *G. syndetika.* while the smallest differentiation was observed between *G. canescens and G. syndetika*. The divergence among the two *Glycine* species in group A was small (Table 2).

**Table 2.** Estimates of evolutionary divergence among *Glycine* species based on 76 DNA sequence analyses of orthologous genes identified from 76 orthologous gene sets that produced a single amplicon in at least six of the eight *Glycine* species analyzed using species specific PCR primers

|  | *G.max* | *G. soja* | *G.*<br>*canescens* | *G.*<br>*cyrtoloba* | *G.*<br>*falcata* | *G.*<br>*stenophita* | *G.*<br>*syndetika* |
| --- | --- | --- | --- | --- | --- | --- | --- |
| *G.max* | 0 |  |  |  |  |  |  |
| *G. soja* | 0.009 | 0 |  |  |  |  |  |
| *G. canescens* | 0.042 | 0.043 | 0 |  |  |  |  |
| *G. cyrtoloba* | 0.050 | 0.049 | 0.041 | 0 |  |  |  |
| *G. falcata* | 0.052 | 0.048 | 0.045 | 0.047 | 0 |  |  |
| *G. stenophita* | 0.042 | 0.042 | 0.032 | 0.037 | 0.045 | 0 |  |
| *G. syndetika* | 0.046 | 0.044 | 0.019 | 0.046 | 0.047 | 0.036 | 0 |
| *G. tomentella*<br>*D3* | 0.043 | 0.041 | 0.025 | 0.041 | 0.042 | 0.032 | 0.028 |

### Trans-specific polymorphism among *Glycine* species

Of the 52 genes, 41 genes contained 1 to 40 trans-specific polymorphic loci. There were ten genes with at least 10 trans-specific polymorphisms (Figure 5). Among the 316 trans-specific polymorphic loci, a total of 152, 81, 61 and 22 were A/G (T/C), A/C (T/G), A/T (T/A), and C/G (G/C), respectively. The transitions (i.e., C/T or A/G) occurred more often than any of the transversions A/C, A/T or C/G.

**Figure 5.**
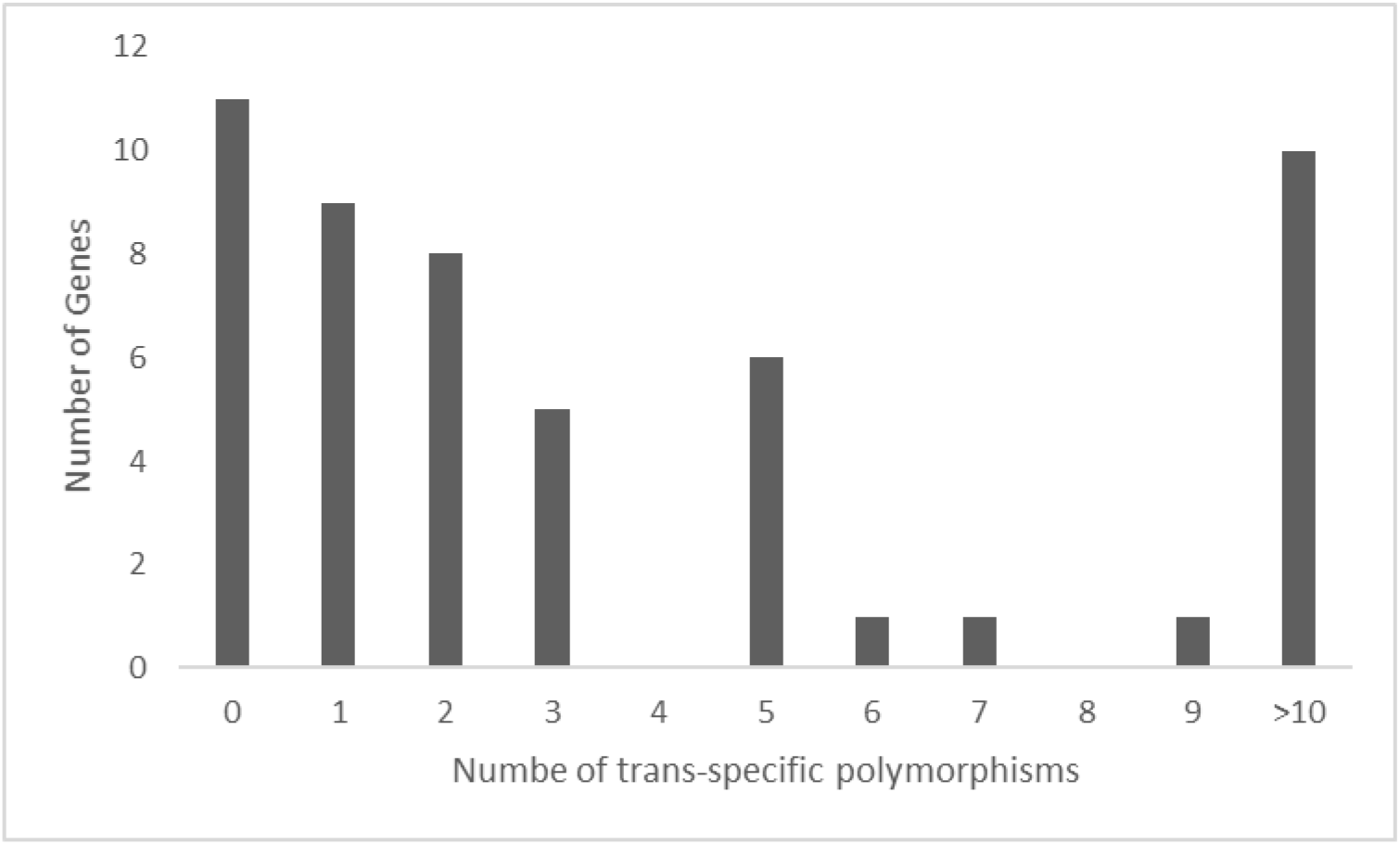
Distribution of genes with different number of trans-specific polymorphisms.

Of the 316 trans-specific polymorphic loci, a total of 223, 66, 9, 14, and 4 were trans-specific among 2, 3, 4, 5 or 6 *Glycine* species, respectively. Further analysis showed that *G. max* vs. *G. syndetika* had the highest number of trans-specific polymorphic loci, followed by *G. syndetika vs. G. tomentella*, *G. falcata* vs. *G. syndetika*, *G. soja vs. G. syndetika, and G. canescens vs. G. max.* Interestingly, *G. syndetika* had the highest number of trans-specific polymorphisms with other species, but *G. cyrtoloba* had the lowest (Table 3). In general, the trans-specific polymorphic loci were more likely observed between the species that were less divergent from each other.

**Table 3.** Number of trans-specific polymorphisms between *Glycine* species

| Species | Number of trans-specific polymorphisms |
| --- | --- |
| *G. max*/*G. syndetika* | 59 |
| *G. syndetika*/*G. tomentella* | 47 |
| *G. falcata*/*G. syndetika* | 44 |
| *G. soja*/*G. syndetika* | 41 |
| *G. canescens*/*G. max* | 37 |
| *G. falcata*/*G. soja* | 36 |
| *G. canescens*/*G. falcata* | 34 |
| *G. canescens*/*G. tomentella* | 34 |
| *G. canescens*/*G. syndetika* | 33 |
| *G. max*/*G. soja* | 32 |
| *G. falcata*/*G. tomentella* | 30 |
| *G. falcata*/*G. max* | 29 |
| *G. max*/*G. tomentella* | 25 |
| *G. soja*/*G. tomentella* | 25 |
| *G. stenophita*/*G. syndetika* | 24 |
| *G. max*/*G. stenophita* | 20 |
| *G. canescens*/*G. soja* | 19 |
| *G. soja*/*G. stenophita* | 17 |
| *G. stenophita*/*G. tomentella* | 16 |
| *G. falcata*/*G. stenophita* | 15 |
| *G. cyrtoloba*/*G. falcata* | 11 |
| *G. cyrtoloba*/*G. max* | 11 |
| *G. cyrtoloba*/*G. syndetika* | 11 |
| *G. canescens*/*G. cyrtoloba* | 10 |
| *G. canescens*/*G. stenophita* | 7 |
| *G. cyrtoloba*/*G. tomentella* | 5 |
| *G. cyrtoloba*/*G. soja* | 1 |
| *G. cyrtoloba*/*G. stenophita* | 1 |
| *G. falcata*/*G. cyrtoloba* | 1 |

### Test of gene for departure from neutrality within species

The Tajima D values for 18-42% of the genes within each species were less than −2, and only a small percentage of the genes (3-17%) had positive D values among species but all were less than 2 (Table 4, Supplementary File 3). The average D value of the genes varied from −1.07 to −1.77 among the species. These results suggested that most genes in each species have been subjected to a recent bout of directional selection due to selective sweep or rapid population expansion.

**Table 4.** Number and percentage of genes with Tajima’s D<-2 and average of Tajima’s D value for genes in each species

| Species | Number<br>of genes<br>with $D < -2$ | Number<br>of genes<br>with $D > -2$ | Percentage<br>of genes<br>with $D < -2$<br>(%) | Average<br>$D$ value |
| --- | --- | --- | --- | --- |
| *G. max* | 10 | 25 | 28.57 | -1.77 |
| *G. soja* | 9 | 18 | 33.33 | -1.38 |
| *G. canescens* | 17 | 33 | 34.00 | -1.68 |
| *G. cyrtoloba* | 17 | 21 | 44.73 | -1.37 |
| *G. falcata* | 12 | 31 | 27.91 | -1.57 |
| *G. stenophita* | 12 | 37 | 24.49 | -1.43 |
| *G. syndetika* | 9 | 42 | 17.64 | -1.07 |
| *G. tomentella* D3 | 23 | 28 | 45.01 | -1.69 |

### HKA test of genes under evolution selection between *G. max* and other *Glycine* species

Because of missing genic sequence and lack of sequence polymorphism in some accessions, HKA values between *G. max* and other species were obtained for 40 of the 52 genes. Of the 169 HKA values acquired between *G. max* and some other *Glycine* species, a total of 23 (13.6%) from 15 genes were significant at P=0.l0, indicating deviation from neutral evolution expectation. The number of genes with significant HKA values between *G. max* and each of other species varied from 2 to 5 (Supplementary File 4). The levels of within- and between-population DNA variation correlated well for most genes (86.4%) between each *Glycine species* and *G. max*.

## DISCUSSION

### Nucleotide diversity

The sets of *Glycine* accessions used in this study were selected to represent the geographical range of the origins of the respective species for the purpose of obtaining a representative sampling in order to provide reliable estimates of genetic diversity. An average genetic diversity of *G. soja* was twice that of *G. max* which was similar to the previous report (Hyten et al. 2006). The values of nucleotide diversity reported by Hyten et al. (2006) were *θ*=0.00235 for 26 *G. soja* accessions collected from China, Korea, Taiwan, Russia and Japan and *θ=* 0.00099 for a set of *G. max* genotypes consisting of 52 Asian landraces, 17 N. American ancestral cultivars and 25 N. American elite cultivars released between 1977 and 1990. Very similar nucleotide diversity values for *G. max* of *θ*=0.00097 and *θ*=0.00099 were reported by Zhu et al. (2003) and Choi et al. (2007), respectively. Of all the perennials, *G. canescens* has the most extensive geographic distribution across Australia (Brown et al. 1990; Gonzalez-Orozco et al. 2012), mostly in very dry areas (Figure 1), while the other perennial species have more localized ranges. As might be expected by its broad geographical distribution and the presence of at least three subgroups within the species as reported by Brown et al. (1990), *G. canescens* had the highest nucleotide diversity of the *Glycine* species which was approximately four times higher than that of *G. max* as estimated by *θ*. With the exception of *G. tomentella* D3 which had unexpectedly low nucleotide diversity (*θ*=0.0016), the other perennial species, *G. cyrtoloba*, *G. falcata*, *G. stenophita*, and *G. syndetika*, had a mean nucleotide diversity (*θ*) higher than that of the annual *G. soja* (*θ*=0.0022) (Table 1). However, the average diversity of the *Glycine* species estimated by *π* was not in complete agreement with the *θ* values. *G. canescens* also showed the highest diversity estimated as *π,* whereas *G. falcata* and *G. stenophita* had a relatively low nucleotide diversity (*π*) as compared with their *θ* values.

There has been no extensive genetic diversity analysis reported on the perennial *Glycine* species because of a lack of nucleotide sequence availability, so it is difficult to compare the current estimation of genetic diversity with other studies. However, we used a total of 63 to 74 orthologous gene sets in the annual and perennial *Glycine* species that were distributed on 19 of the 20 soybean chromosomes (Figure 2). This is the first report to provide an estimate of genetic diversity of the *Glycine* species based upon sequences distributed across the genomes of the annual species and given the close relationship of the annual and perennial species, these sequences are also very likely to be spread across the genomes of the perennial species.

### Phylogenetic analysis

An important step after the collection of new germplasm is to determine its relationship with other known species by comparing morphological traits, by cytological and crossing studies and/or by the analysis of molecular variability. The early phylogenetic analyses of the perennial *Glycine* species were conducted mostly using interspecies crossability and cytogenetic studies with the ultimate intention of the discovery and introgression of beneficial genes/alleles into cultivated soybeans. These efforts resulted in the construction of the first nuclear genome designation of the genus *Glycine* composed of six diploid genome groups (Singh and Hymowitz 1985a) which ultimately included nine genome groups from A to I (Hymowitz et al. 1998) including 25 perennial species (Doyle et al. 2004; Sherman-Broyles et al. 2014). Based upon these previous analyses, the eight *Glycine* species used in the current study belong to six of the nine genome groups. The two annual *Glycine* species, *G. soja* and *G. max*, belong to the G genome group; the perennial species *G. canescens* and *G. syndetika* belong to the A group; *G. stenophita* to the group B’ which was integrated into the genome grouping system by Brown *et al*. (2002); *G. cyrtoloba* to the C group; *G. tomentella* D3 to the D group, and *G. falcata* to the F group. There have been studies reported of the phylogenetic relationships of the *Glycine* species based on nucleotide sequence variation in a single or small number of nuclear DNA sequences (Kollipara et al. 1997; Doyle et al. 1999a; Brown et al. 2002) or chloroplast DNA sequence (Doyle et al. 1990; Sakai et al. 2003). As might be expected, the tree estimations from different studies were different (Doyle et al. 2003b) because different genes had a different gene history which might not represent the history of the taxa (Rokas et al. 2003). In this study, we used approximately 87.6 Kbp of common sequence from 52 orthologous gene sets present in the 77 accessions of the eight *Glycine* species and two out-groups, *Phaseolus vulgaris* L. and *Vigna unguiculata* (L.) Walp, for the estimation of the phylogenetic relationships. This analysis should provide a reasonable estimation of the phylogenetic relationships of the eight *Glycine* species.

The phylogenetic relationships estimated in this study agreed with a sister relationship between the monophyletic annual and perennial subgenera reported in the previous studies (Doyle et al. 2003a) and it agreed with the relationships based upon Histone H3-D (Sherman-Broyles et al. 2014), except that *G. cyrtoloba* of the C genome group and *G. stenophita* of the B’ group formed a sister clade (Figure 3). The close relationship between *G. cyrtoloba* and the B genome group was suggested by Singh (2016). Based upon a cytogenetic analysis, Singh also reported that the D genome group had evolved from the A group which agreed with the phylogenetic analysis performed in this study where the A group species, *G. canescens* and *G. syndetika* and the D group species, *G. tomentella*, formed a sister clade (Figure 3).

Of the six perennial species, *G. canescens* is the species with the most geographically widespread distribution in Australia (Brown et al. 1990; Gonzalez-Orozco et al. 2012). Brown *et al*. (1990) estimated the genetic structure of *G. canescens* using the variation of 11 isozymes and reported that *G. canescens* could be classified into five groups, the groups G1, G2, and G3 which had different isozyme variation patterns, the U group which did not belong to any of the other groups but shared more alleles with group G3 and the X group which was morphologically similar to *G. clandestina*. In addition, there were *G. canescens* accessions collected more recently whose genome groups have not been determined. Two of the *G. canescens* accessions used in this study belong in this latter category. In Figure 3 the clade of *G. canescens* includes only 9 accessions because a number of the 52 orthologous gene sets used in the analysis were not present in the two *G. canescens* accessions PI505160 and PI591574. To confirm the relationships among the 9 *G. canescens* accessions we performed a separate phylogenetic analysis using 367 orthologous gene sets in 11 accessions which added two more group G1 accessions to the analysis. In both analyses the genome group G1 accessions were clustered in as single clade as were the group G3 accessions (Figure 3 and Figure 4A). In figure 4B, the group G1 accessions originated from the Northern Territory, South Australia, New South Wales, and Victoria, however, their geographic origins were limited to longitudes between 134°E and 143°E and latitudes between 25°S and 35°S where the maximum physical distance among the geographical origins of the four G1 accessions was 1,270 Km followed by 946 Km and 684 Km (Figure 4B). The group G3 accessions were from New South Wales, South Australia, and Queensland where the average precipitation is greater than the areas from which the other accessions originated (Figure 1) and as was the case for the group G1 accessions, as the distance between the geographic origins of the accessions increased their phylogenic relationships were more distant. (Figure 4B). The two ND and U accessions were in a distinct clade in the phylogenetic tree using the 367 gene sets (Figure 4A) whereas one of ND accessions was clustered with the group G1 accessions in the tree using the 52 gene sets (Figure 3). The ND and U accessions originated from Western Australia which is geographically apart from the regions where the group G1, G2, and G3 accessions originated (Figure 4B). Based on the geographic origins of the ND and U accessions, the clustering using the 367 gene sets would appear to be more reliable. In this study we used only one group G2 accession from the Northern Territory so that we do not know the relationships among the group G2 accessions. However, it was assumed that the genome group G2 was more closely related to the group G1 than the group G3 based upon the phylogenetic tree in Figure 4A. Based on the discussion above, in the case of *G. canescens,* the phylogenetic analysis using the 367 gene sets was more reliable and agreed with the groupings as reported by Brown *et al*. (1990) with the high bootstrap values rather than the phylogenetic tree estimated using the 52 gene sets. Clearly, *G. canescens* appeared to be the most diverse of the perennial species examined in this study based upon sequence diversity, geographical origins and the phylogenetic analysis.

This study is the first to provide an estimate of genetic diversity of the perennial *Glycine* species and we found that all the perennial species had higher genetic diversity (*θ* and *π*) than the annual species, except *G. tomentella* D3 and *G. falcata*, based upon the analysis of a total of 63 - 74 orthologous gene sets. The greater genetic diversity of most of the perennial *Glycine* offers the hope of variation for genetically controlled traits that is not present in the annual *Glycine* species. Some of the perennial *Glycine* have been adapted to the environments were the average annual precipitation on the Australian continent was approximately 450 mm for the last 100 years (http://www.abs.gov.au). The regions where *G. canescens* are found are the driest and hottest areas in Australia with an annual precipitation of less than 300 mm and an average daily maximum temperature of 33 ͦ C in January. All of the other perennial *Glycine* are geographically localized from the northeastern to southeastern coastal areas in Australia (Figure 1) where the average maximum temperature in January is 27 ͦ C with relatively higher annual precipitation ranging from 500 to 1600 mm. Therefore, we can assume that the perennial *Glycine,* and particularly *G. canescens* with its origin in the driest and warmest regions of Australia, could have numerous stress related genes/alleles to survive in drought and high temperatures which would be the candidate traits that need to be integrated into cultivated soybean to cope with future climate change. Given the likelihood of climate change, it seems likely that genes controlling resistance to heat and drought will be of particular importance in future soybean cultivars. Since heat and drought stress resistance are likely to be controlled by a number of genes, and given the rapidity with which high density genetic maps can now be developed using genotyping by sequencing (Elshire et al. 2011), gene identification and cloning from the perennial *Glycine* species may become a relatively rapid process that can be used as a source of genes/alleles in cultivated soybean.

### Divergence analysis among *Glycine* species

Among the perennial species, the three species *G. tomentella*, *G. canescens, and G. syndetika* were less divergent from each other, but were the most divergent from *G. falcata* based on the Tajima-Nei distance analysis method. The estimates of divergence among species based on the Tajima-Nei distance analysis were consistent with those based on other models such as p-distance (Nei and Kumar 2000), Jukes-Cantor distance (Jukes and Cantor 1969), the Kimura 2-parameter distance (Kimura 1980) and the Tamura-Nei distance (Tamura and Nei 1993). The relationship among *Glycine* species based on sequence divergence was also consistent with that inferred by the phylogenic tree (Figure 3) and the outcome of hybridization among *Glycine* species. We observed that the *G. max* was the most divergent from *G. falcata,* followed by *G. cyrtoloba*, *G. syndetika*, *G. tomentella* D3, *G. stenophita* and *G. canescens*, this conclusion was consistent with the crossability report of soybean cultivars ‘Lincoln’ and ‘Hark’ with *G. tomentella*, *G. canescens, G. falcata, G. tabacina, G. latrobeana, G. clandestina, and G. latifolia*, respectively (Broue et al. 1982). From those crosses, hybrid seeds were only obtained from the crosses with *G. tomentella* and *G. canescens*. Successful hybridization of cultivated soybean with *G. tomentella* and *G. canescens* was also reported in a number of other studies (Newell et al. 1987; Newell and Hymowitz 1982; Singh and Hymowitz 1985b; BodaneseZanettini et al. 1996). Besides cytogenetic differences between *G. falcata* and other *Glycine species* reported by Putievsky and Broue (1979), the habit of pod production between *G. falcata* and other *Glycine* species was also different. Only *G. falcata* develops pods underground like peanut. The relationship among *Glycine* species and the estimates of divergence between soybean and the other *Glycine* species provided important information for selection of perennial species that will most likely be successful to hybridize with cultivated soybean.

### Trans-specific mutations

The higher occurrence of transitions (i.e., C/T or A/G) than expected by chance and transversions were consistent with the results reported by Brown et al. (1982) and Keller et al. (2007). Transition mutations are more easily generated due to molecular shape, and are less likely to be removed by natural selection because they more often create synonymous amino acids as demonstrated in influenza virus and HIV by Lyons and Lauring (2017). Although trans-specific polymorphic loci were more frequently observed between species that were less divergent, we didn’t identify a high number of those loci between *G. max* and *G. soja.* Because the sequence of 52 genes was highly conserved among *Glycine species*, most sequences that were polymorphic in perennial species were fixed within *Glycine soja* and *G. max*, thus resulting in a limited number of polymorphic loci. In addition, we observed that the same trans-specific mutations frequently occurred in more than three species in the same gene, which suggested that *Glycine* species may have experienced very similar evolutionary selection or the *Glycine* species may share the same mechanism to adapt to their changing environments.

Our results showed that *G. max* shared the highest number of trans-specific polymorphism with *G. syndetika*, followed by *G. canescens*. The results are not unexpected, previous studies of the perennial species classified *G. syndetika* and *G. canescens* into genome group A based on interspecific crossability, meiotic chromosome pairing or seed protein electrophoresis (Hymowitz et al. 1998; Sherman-Broyles et al. 2014; Singh and Hymowitz 1985a) and the two species were sister clade of *G. tomentella* (D genome group) (Doyle et al. 2003a; Sherman-Broyles et al. 2014). Broue et al. (1982) crossed the *G. max* with 7 perennial species, only the mating with *G. tomentella and G. canescens, the* two closest relatives of *G. syndetika,* was successful. Although no attempt has been made to cross *G. syndetika* with *G. max*, the crossability of *G. tomentella and G. canescens* with *G. max* implies that the *G. syndetika* is not divergent from *G. max*. Our divergence analysis also suggested that the *G. max* was less divergent from *G. syndetika* than *G. falcata and G. cyrtoloba* (Table 2).

A number of studies have reported the relationship among the *Glycine* species (Kollipara et al. 1997; Doyle et al. 1990; Doyle et al. 1999a; Sherman-Broyles et al. 2014), and conclusions from those studies generally varied depending on taxon sampling and the locus that was used (Sherman-Broyles et al. 2014). Of these studies, only a limited number included both *G. max* and *G. syndetika*, Sherman-Broyles et al. (2014) analyzed the pair-wise SNPs among *Glycine* species plastomes and concluded that the distance of *G. max* vs. *G. syndetika* was smaller than *G.max vs. G. falcata*, and *G. max vs. G. cyrtoloba.* In another study involving wild soybean (*G. soja*), the closest relative of *G. max*, and *G. canescen,* the closest relative of *G. syndetika*, was lower than *G. soja vs. G. falcata*, and *G. soja vs. G. cyrtoloba* based on restriction-endonuclease site variation of chloroplast DNA (Doyle et al. 1990). Because species topology may change according to the order of the species arranged on the phylogenic tree, topology sometime may not represent the true divergence among species, thus divergence estimates by distance measures are more reliable.

### Neutral selection of genes

We observed that most genes had a negative Tajima’s D value, and approximately 17-45% of the genes among *Glycine* species experienced significant adaptive evolution. The dominance of genes with negative D value was consistent with our observation of a high *ϴ vs.* low *π* statistics of genes in each species (Table 1). The significant and negative D value for the majority of the genes in *Glycine* species resulting from an overall excess of rare polymorphisms is strong evidence of the recent population expansion or selective sweep (Akey et al. 2004). Significant and negative Tajima’s D values were also reported in previous studies for most genes in human (Stephens et al. 2001) and *Drosophila melanogaster* populations (Haddrill et al. 2005). Of the 313 genes analyzed in a human population with 82 unrelated individuals of diverse ancestry, a total of 281 (89.8%) showed a negative Tajima’s D value with an average of −0.97 (Stephens et al. 2001).

Approximately 14% of the genes between *G. max* and other *Glycine* species deviated from neutral evolution based on the HKA test, the percentage was lower than that reported between *Drosophila simulans and Drosophila meleanogaster* and between humans and monkeys, e.g. approximately 50% of the amino acid substitutions within genes between *Drosophila simulans and Drosophila meleanogaster* were driven by positive selection (Brookfield and Sharp 1994; Sawyer and Hartl 1992), and approximately 35% between humans and old-world monkeys experienced adaptive evolution (Fay et al. 2001). Genes with negative significant Tajima’s D but insignificant HKA values were previously reported in *Drosophila melanogaster* populations by Haddrill et al. (2005). Although 17-45% gene sequence had negative significant Tajima’s D, our data showed that most gene sequences were homogeneous in levels of polymorphism and divergence between *G. max* and other *Glycine* species, indicating that all the *Glycine* perennial species may have experienced very similar evolutionary selection as inferred by the trans-specific mutation analysis.

Previous reports showed that missing data or sequence errors may cause the bias of Tajima’s D values. Korneliussen et al. (2013) observed that Tajima’s *D* calculated using genotypes called from next generation sequence data could lead to biased results and the level of bias depends on the sequencing depths and error rates. Ferretti et al. (2012) reported that missing data can also contribute to the bias. In this study, we only observed a small percentage (2%) of the genes that had a Tajima’s D value of less than −3 among 8 species when site sequence coverage was above 70%. And above the 70% threshold, the Tajima’s D estimates were generally stable for almost all genes. However, when the site sequence coverage was lowered or the option of pairwise deletion of the sequence with missing in Mega program was used, the percentage of genes with biased Tajima’s D value increased. This is because the length of assembled sequence for each gene varied among accessions within each species, only very few accessions may have sequences at the beginning or/and end of the gene sequence alignment. The small number of sequence dramatically reduced the *π* value but relatively has a small impact on *ϴ*, thus cause a D bias toward a small negative value. We therefore suggest a high threshold of site coverage for the analysis. The report of Tajima’s D value in perennial *Glycine* species was limited. In soybean, the range of genome-wide Tajima’s D values was from −3 to 4 based on the sequence analysis of 24 accessions (Lam et al, 2010), our results generally were consistent with this study. We also observed 17-45% of the genes among species with significant negative Tajima D values in this study, the positive selection of genes in plant is common, e.g. in common bean, Schmutz et al (2014) identified 1,835 Mesoamerican and 748 Andean candidate genes associated with domestication and all candidates had a negative Tajima’s *D* value.

The sequence random error may also cause Tajima’s D bias, however, the error may exist but the error rate is likely low in this study. The conclusion is supported by the facts that the rate of SNPs with more than two alleles *vs.* all SNPs is low ranging from 0.15% to 1.63% among *Glycine* species; the SNP alleles in one species generally matched one base or both bases (if polymorphic in any other species) in other *Glycine* species.

## AUTHOR CONTRIBUTION STATEMENT

P.B.C. and Q.S provided project planning and coordination. E.Y.H, C.V.Q. and E.W.F. performed RNA extraction, library construction, sequencing preparation and PCR amplification. S.G.S., C.V.Q, E.W.F. and P.E. performed sequence analysis. E.Y.H, Q.S., H.W, S.A, L.C, M.A.F and P.B.C performed sequence data analyses and interpretation. E.Y.H, Q.S and P.B.C. prepared the manuscript.

## ACKNOWLEDGEMENTS

This work is part of the SoyMap2 project and was funded by National Science Foundation Grant 0822258. We thank Rob Parry and Chris Pooley for their technical support in assembling the necessary hardware and software required for the data analysis

## LITERATURE CITED

1. Akey, J.M., M.A. Eberle, M.J. Rieder, C.S. Carlson, M.D. Shriver et al., 2004 Population history and natural selection shape patterns of genetic variation in 132 genes. PLoS biology 2 (10):e286.

2. Barker, D.G., T. Pfaff, D. Moreau, E. Groves, S. Ruffel et al., 2006 Growing M. truncatula: choice of substrates and growth conditions. Medicago truncatula handbook. Retrieved from https://www.noble.org/Global/medicagohandbook/pdf/GrowingMedicagotruncatula.pdf.

3. Bauer, S., T. Hymowitz, and G. Noel, 2007 Soybean cyst nematode resistance derived from Glycine tomentella in amphiploid (G. max X G. tomentella) hybrid lines. Nematropica 37 (2):277–286.

4. BodaneseZanettini, M.H., M.S. Lauxen, S.N.C. Richter, S. CavalliMolina, C.E. Lange et al., 1996 Wide hybridization between Brazilian soybean cultivars and wild perennial relatives. Theoretical and Applied Genetics 93 (5-6):703–709.

5. Brookfield, J.F.Y., and P.M. Sharp, 1994 Neutralism and Selectionism Face up to DNA Data. Trends in Genetics 10 (4):109–111.

6. Broue, P., J. Douglass, J.P. Grace, and D.R. Marshall, 1982 Interspecific Hybridization of Soybeans and Perennial Glycine Species Indigenous to Australia Via Embryo Culture. Euphytica 31 (3):715–724.

7. Brown, A.H.D., J.J. Burdon, and J.P. Grance, 1990 Genetic structure of *Glycine canescens*, a perennial relative of soybean. Theor. Appl. Genet. 79:729–736.

8. Brown, A.H.D., J.L. Doyle, J.P. Grace, and J.J. Doyle, 2002 Molecular phylogenetic relationships within and among diploid races of *Glycine tomentella* (Leguminosae). Aust Syst Bot 15:37–47.

9. Brown, W.M., E.M. Prager, A. Wang, and A.C. Wilson, 1982 Mitochondrial DNA sequences of primates: tempo and mode of evolution. Journal of Molecular Evolution 18 (4):225–239.

10. Burdon, J.J., 1988 Major Gene Resistance to Phakopsora-Pachyrhizi in Glycine-Canescens, a Wild Relative of Soybean. Theoretical and Applied Genetics 75 (6):923–928.

11. Carlson, C.S., D.J. Thomas, M.A. Eberle, J.E. Swanson, R.J. Livingston et al., 2005 Genomic regions exhibiting positive selection identified from dense genotype data. Genome research 15 (11):1553–1565.

12. Choi, I.Y., D.L. Hyten, L.K. Matukumalli, Q.J. Song, J.M. Chaky et al., 2007 A soybean transcript map: Gene distribution, haplotype and single-nucleotide polymorphism analysis. Genetics 176 (1):685–696.

13. Craig, D.W., J.V. Pearson, S. Szelinger, A. Sekar, M. Redman et al., 2008 Identification of genetic variants using bar-coded multiplexed sequencing. Nat. Methods 5 (10):887–893.

14. Darriba, D., G.L. Taboada, R. Doallo, and D. Posada, 2012 jModelTest 2: more models, new heuristics and parallel computing. Nat Methods 9 (8):772.

15. Doyle, J., J. Doyle, and A. Brown, 1990 A chloroplast DNA phylogeny of the wild perennial relatives of soybean (*Glycine* subgenus *Glycine*): congruence with morphological and crossing groups. Evolution 44:371–389.

16. Doyle, J.J., J. Doyle, L., J. Rauscher, T., and A.H.D. Brown, 2004 Evolution of the perennial soybean polyploid complex (*Glycine* subgenus *Glycine*): a study of contrast. Biol. J. Linn. Soc. Lond. 82:583–597.

17. Doyle, J.J., J.L. Doyle, and A.H. Brown, 1999a Incongruence in the diploid B-genome species complex of Glycine (Leguminosae) revisited: histone H3-D alleles versus chloroplast haplotypes. Mol Biol Evol 16 (3):354–362.

18. Doyle, J.J., J.L. Doyle, and A.H. Brown, 1999b Origins, colonization, and lineage recombination in a widespread perennial soybean polyploid complex. Proc. Natl. Acad. Sci. U S A 96 (19):10741–10745.

19. Doyle, J.J., J.L. Doyle, and C. Harbison, 2003a Chloroplast-expressed glutamine synthetase in Glycine and related Leguminosae: phylogeny, gene duplication, and ancient polyploidy. Systematic botany. v. 28:567–577.

20. Doyle, J.J., J.L. Doyle, J.T. Rauscher, and A.H.D. Brown, 2003b Diploid and polyploid reticulate evolution throughout the history of the perennial soybeans (*Glycine* subgenus *Glycine*). New Phytologist 161:121–132.

21. Doyle, J.J., V. Kanazin, and R.C. Shoemaker, 1996 Phylogenetic utility of histone H3 intron sequences in the perennial relatives of soybean (Glycine: Leguminosae). Molecular Phylogenetics and Evolution 6 (3):438–447.

22. Edgar, R.C., 2004 MUSCLE: multiple sequence alignment with high accuracy and high throughput. Nucleic Acids Res. 32 (5):1792–1797.

23. Edgar, R.C., 2010 Search and clustering orders of magnitude faster than BLAST. Bioinformatics 26 (19):2460–2461.

24. Egan, A.N., and J. Doyle, 2010 A comparison of global, gene-specific, and relaxed clock methods in a comparative genomics framework: dating the polyploid history of soybean (Glycine max). Syst Biol 59 (5):534–547.

25. Elshire, R.J., J.C. Glaubitz, Q. Sun, J.A. Poland, K. Kawamoto et al., 2011 A Robust, Simple Genotyping-by-Sequencing (GBS) Approach for High Diversity Species. Plos One 6 (5).

26. Ewing, B., L. Hillier, M.C. Wendl, and P. Green, 1998 Base-calling of automated sequencer traces using phred. I. Accuracy assessment. Genome Res 8 (3):175–185.

27. Fay, J.C., G.J. Wyckoff, and C.I. Wu, 2001 Positive and negative selection on the human genome. Genetics 158 (3):1227–1234.

28. Ferretti, L., E. Raineri, and S. Ramos-Onsins, 2012 Neutrality tests for sequences with missing data. Genetics 191 (4):1397–1401.

29. Gonzalez-Orozco, C.E., A.H.D. Brown, N. Knerr, J.T. Miller, and J.J. Doyle, 2012 Hotspots of diversity of wild Australian soybean relatives and their conservation in situ. Conserv Genet 13:1269–1281.

30. Grant, J.E., A.H.D. Brown, and J.P. Grace, 1984 Cytological and isozyme diversity in Glycine tomentella Hayata (Leguminosae). Australian journal of botany:665–677.

31. Guindon, S., J.F. Dufayard, V. Lefort, M. Anisimova, W. Hordijk et al., 2010 New algorithms and methods to estimate maximum-likelihood phylogenies: assessing the performance of PhyML 3.0. Syst Biol 59 (3):307–321.

32. Haddrill, P.R., K.R. Thornton, B. Charlesworth, and P. Andolfatto, 2005 Multilocus patterns of nucleotide variability and the demographic and selection history of Drosophila melanogaster populations. Genome research 15 (6):790–799.

33. Hamady, M., J.J. Walker, J.K. Harris, N.J. Gold, and R. Knight, 2008 Error-correcting barcoded primers for pyrosequencing hundreds of samples in multiplex. Nat. Methods 5 (3):235–237.

34. Hartman, G.L., T.C. Wang, and T. Hymowitz, 1992 Sources of Resistance to Soybean Rust in Perennial Glycine Species. Plant Disease 76 (4):396–399.

35. Hudson, R.R., M. Kreitman, and M. Aguadé, 1987 A test of neutral molecular evolution based on nucleotide data. Genetics 116 (1):153–159.

36. Hymowitz, T., R.J. Singh, and K.P. Kollipara, 1998 The genomes of the *Glycine*. Plant Breeding Reviews 16:289–317.

37. Hyten, D.L., Q.J. Song, Y.L. Zhu, I.Y. Choi, R.L. Nelson et al., 2006 Impacts of genetic bottlenecks on soybean genome diversity. Proceedings of the National Academy of Sciences of the United States of America 103 (45):16666–16671.

38. Innes, R.W., C. Ameline-Torregrosa, T. Ashfield, E. Cannon, S.B. Cannon et al., 2008 Differential accumulation of retroelements and diversification of NB-LRR disease resistance genes in duplicated regions following polyploidy in the ancestor of soybean. Plant Physiol 148 (4):1740–1759.

39. Jukes, T.H., and C.R. Cantor, 1969 Evolution of protein molecules. In Munro HN, editor, Mammalian Protein Metabolism, pp. 21–132. Academic Press, New York.

40. Keim, P., T.C. Olson, and R.C. Shoemaker, 1988 A rapid protocol for isolating soybean DNA. Soybean Genet. Newsl. 15:150–152.

41. Keller, I., D. Bensasson, and R.A. Nichols, 2007 Transition-transversion bias is not universal: a counter example from grasshopper pseudogenes. PLoS genetics 3 (2):e22.

42. Kimura, M., 1980 A Simple Method for Estimating Evolutionary Rates of Base Substitutions through Comparative Studies of Nucleotide-Sequences. Journal of Molecular Evolution 16 (2):111–120.

43. Kollipara, K.P., R.J. Singh, and T. Hymowitz, 1994 Genomic diversity and multiple origins of tetraploid (2n = 78, 80) Glycine tomentella. Genome 37 (3):448–459.

44. Kollipara, K.P., R.J. Singh, and T. Hymowitz, 1997 Phylogenetic and genomic relationships in the genus Glycine Willd. based on sequences from the ITS region of nuclear rDNA. Genome 40 (1):57–68.

45. Korneliussen, T.S., I. Moltke, A. Albrechtsen, and R. Nielsen, 2013 Calculation of Tajima’s D and other neutrality test statistics from low depth next-generation sequencing data. BMC Bioinformatics 14 (1):289.

46. Kumar, S., G. Stecher, and K. Tamura, 2016 MEGA7: Molecular Evolutionary Genetics Analysis Version 7.0 for Bigger Datasets. Molecular biology and evolution 33 (7):1870–1874.

47. Lechner, M., S. Findeiss, L. Steiner, M. Marz, P.F. Stadler et al., 2011 Proteinortho: detection of (co-)orthologs in large-scale analysis. BMC Bioinformatics 12:124.

48. Lam, H.M., Xu, X., Liu, X., Chen, W., Yang, G., et al., 2010 Resequencing of 31 wild and cultivated soybean genomes identifies patterns of genetic diversity and selection. Nature genetics, 42(12) p1053.

49. Lyons, D.M., and A.S. Lauring, 2017 Evidence for the selective basis of transition-to-transversion substitution bias in two RNA viruses. Molecular biology and evolution 34 (12):3205–3215.

50. Nei, M., and S. Kumar, 2000 Molecualr Evoluation and Phylogenetics. Oxford University Press, New York.

51. Newell, C.A., X. Delannay, and M.E. Edge, 1987 Interspecific Hybrids between the Soybean and Wild Perennial Relatives. Journal of Heredity 78 (5):301–306.

52. Newell, C.A., and T. Hymowitz, 1982 Successful Wide Hybridization between the Soybean and a Wild Perennial Relative, G-Tomentella Hayata. Crop Science 22 (5):1062–1065.

53. Parks, D.H., T. Mankowski, S. Zangooei, M.S. Porter, D.G. Armanini et al., 2013 GenGIS 2: Geospatial Analysis of Traditional and Genetic Biodiversity, with New Gradient Algorithms and an Extensible Plugin Framework. Plos One 8 (7).

54. Putievsky, E., and P. Broue, 1979 Cytogenetics of Hybrids among Perennial Species of Glycine Subgenus Glycine. Australian journal of botany 27 (6):713–723.

55. Ratnaparkhe, M.B., R.j. Singh, and J.J. Doyle, 2011 Glycine. In C. Kole, Wild Crop Relatives: Genomic and Breeding Resources: Legume Crops and Forages (pp. 83–116). Springer-Verlag Berlin Heidelberg.

56. Riggs, R.D., S. Wang, R.J. Singh, and T. Hymowitz, 1998 Possible transfer of resistance to *Heterodera glycines* from *Glycine tomentella* to soybean. J Nematol 30:547–552.

57. Rokas, A., B.L. Williams, N. King, and S.B. Carroll, 2003 Genome-scale approaches to resolving incongruence in molecular phylogenies. Nature (London, England) Nature. v. 425:798–804.

58. Rozen, S., and H. Skaletsky, 2000 Primer3 on the WWW for general users and for biologist programmers. Methods Mol. Biol. 132:365–386.

59. Sakai, M., K. A., A. Fujii, F.S. Thseng, J. Abe et al., 2003 Phlogenetic relationships of the chloroplast genomes in the genus *Glycine* inferred from four intergenic spacer sequences. Plant Syst. Evol. 239:29–54.

60. Sawyer, S.A., and D.L. Hartl, 1992 Population-Genetics of Polymorphism and Divergence. Genetics 132 (4):1161–1176.

61. Schmutz, J., S.B. Cannon, J. Schlueter, J. Ma, T. Mitros et al., 2010 Genome sequence of the palaeopolyploid soybean. Nature 463 (7278):178–183.

62. Sherman-Broyles, S., A. Bombarely, J. Grimwood, J. Schmutz, and J. Doyle, 2014 Complete plastome sequences from Glycine syndetika and six additional perennial wild relatives of soybean. G3 (Bethesda) 4 (10):2023–2033.

63. Sherman-Broyles, S., A. Bombarely, A.F. Powell, J.L. Doyle, A.N. Egan et al., 2014 The wild side of a major crop: Soybean’s perennial cousins from Down Under. American journal of botany 101 (10):1651–1665.

64. Singh, R.J., 2016 Plant cytogenetics: CRC press.

65. Singh, R.J., and T. Hymowitz, 1985a The genomic relationships among six wild perennial species of the genus *Glycine* subgenus *Glycine* Willd. Theor. Appl. Genet. 71:221–230.

66. Singh, R.J., and T. Hymowitz, 1985b An Intersubgeneric Hybrid between Glycine-Tomentella Hayata and the Soybean, Glycine-Max (L) Merr. Euphytica 34 (1):187–192.

67. Singh, R.J., K.P. Kollipara, and T. Hymowitz, 1992 Genomic relationships among diploid wild perennial species of the genus Glycine Willd. subgenus Glycine revealed by crossability, meiotic chromosome pairing and seed protein electrophoresis. Theor. Appl. Genet. 85:276–282.

68. Singh, R.J., K.P. Kollipara, and T. Hymowitz, 1998 The genomes of *Glycine canescens* F.J. Herm., amd G. tomentella Hayata of Western Australia and their phylogenetic relationships in the genus Glycine Willd. Genome 41:669–679.

69. Stephens, J.C., J.A. Schneider, D.A. Tanguay, J. Choi, T. Acharya et al., 2001 Haplotype variation and linkage disequilibrium in 313 human genes. Science.

70. Tajima, F., 1983 Evolutionary relationship of DNA sequences in finite populations. Genetics 105 (2):437–460.

71. Tajima, F., 1989 Statistical method for testing the neutral mutation hypothesis by DNA polymorphism. Genetics 123 (3):585–595.

72. Tajima, F., and M. Nei, 1984 Estimation of Evolutionary Distance between Nucleotide-Sequences. Molecular biology and evolution 1 (3):269–285.

73. Tamura, K., and M. Nei, 1993 Estimation of the Number of Nucleotide Substitutions in the Control Region of Mitochondrial-DNA in Humans and Chimpanzees. Molecular biology and evolution 10 (3):512–526.

74. Watterson, G.A., 1975 On the number of segregating sites in genetical models without recombination. Theor. Popul. Biol. 7 (2):256–276.

75. Zerbino, D.R., and E. Birney, 2008 Velvet: algorithms for de novo short read assembly using de Bruijn graphs. Genome Res. 18 (5):821–829.

76. Zhu, Y., Q. Song, D. Hyten, C. Van Tassell, L. Matukumalli et al., 2003 Single-nucleotide polymorphisms in soybean. Genetics 163 (3):1123–1134.

77. Zhulidov, P.A., E.A. Bogdanova, A.S. Shcheglov, L.L. Vagner, G.L. Khaspekov et al., 2004 Simple cDNA normalization using kamchatka crab duplex-specific nuclease. Nucleic Acids Res. 32 (3):e37.

